# Using Barcoded Zika Virus to Assess Virus Population Structure *in vitro* and in *Aedes aegypti* Mosquitoes

**DOI:** 10.1101/222984

**Authors:** James Weger-Lucarelli, Selene M. Garcia, Claudia Rückert, Alex Byas, Shelby L. O’Connor, Matthew T. Aliota, Thomas C. Friedrich, David H. O’Connor, Gregory D. Ebel

## Abstract

Arboviruses such as Zika virus (ZIKV, *Flaviviridae; Flavivirus*) replicate in both mammalian and insect hosts where they encounter a variety of distinct host defenses. To overcome these pressures, arboviruses exist as diverse populations of distinct genomes. However, transmission between hosts and replication within hosts can involve genetic bottlenecks, during which population size and viral diversity may be significantly reduced, potentially resulting in large fitness losses. Understanding the points at which bottlenecks exist during arbovirus transmission is critical to identifying targets for preventing transmission. To study these bottleneck effects, we constructed 4 “barcoded” ZIKV clones - 2 with an 8-base-pair degenerate insertion in the 3’ UTR and 2 with 8 or 9 degenerate synonymous changes in the coding sequence, theoretically containing thousands of variants each. We passaged these viruses 3 times each in 2 mammalian and 2 mosquito cell lines and characterized selection of the “barcode” populations using deep sequencing. Additionally, the viruses were used to feed three recently field-caught populations of *Aedes aegypti* mosquitoes to assess bottlenecks in a natural host. The barcoded viruses replicated well in multiple cell lines *in vitro* and *in vivo* in mosquitoes and could be characterized using next-generation sequencing. The stochastic nature of mosquito transmission was clearly shown by tracking individual barcodes in *Ae. aegypti* mosquitoes. Barcoded viruses provide an efficient method to examine bottlenecks during virus infection.

**AUTHOR SUMMARY:** In general, mosquito-borne viruses like ZIKV must replicate in two very different host environments: an insect and a mammalian host. RNA viruses such as ZIKV must maintain genetic diversity in order to adapt to these changing conditions. During this transmission cycle, several barriers exist which can severely restrict viral genetic diversity, causing bottlenecks in the virus population. It is critical to understand these bottlenecks during virus transmission as this will provide important insights into the selective forces shaping arbovirus evolution within and between hots. Here, we employ a set of barcoded ZIKV constructs containing a degenerate stretch of nucleotides that can be tracked using next-generation sequencing. We found that the insertion site in the genome was an important determinant of the resulting diversity of the genetic barcode. We also found that bottlenecks varied between different mosquito populations and patterns of genetic diversity were distinct among individual mosquitoes within a single population, highlighting the randomness of virus dissemination in mosquitoes. Our study characterizes a new tool for tracking bottlenecks during virus transmission *in vivo* and highlights the importance of both viral and host factors on the maintenance of viral diversity.

## INTRODUCTION

Zika virus (ZIKV, *Flaviviridae; Flavivirus*) was responsible for a pandemic first noted on the Pacific Island of Yap in 2007 that expanded throughout the Americas beginning in 2013-14 (1, 2). ZIKV produces either an asymptomatic infection or mild febrile disease in most infected humans. However, the ongoing outbreak has been characterized by a high incidence of malformations in developing fetuses and neurological complications in adults resulting in massive public concern (3, 4). In addition to mosquito-borne transmission, ZIKV is capable of being spread sexually (5), via direct contact (6), and vertically in both humans and mosquitoes (7–9). Understanding the dynamics of transmission is essential for developing strategies to prevent future pandemics caused by ZIKV or other arboviruses.

Transmission bottlenecks are a major stochastic force in virus evolution. In particular, arthropod-borne viruses (arboviruses) must replicate efficiently in two vastly different hosts in order to perpetuate in nature. Several bottlenecks may exist within each host. For example, for mosquito transmission to occur, arboviruses must first infect and replicate within the insect’s midgut epithelial cells and disseminate to secondary tissues such as the fat body, hemocytes, and the nervous system (10). Following infection of these tissues, the virus must then productively infect the salivary glands to eventually be expectorated during feeding. Many of these steps impose genetic bottlenecks that have been shown to rapidly reduce the genetic diversity of arboviruses including West Nile virus (WNV) (11), Powassan virus (12), and Venezuelan equine encephalitis virus (13). Recovery of lost genetic diversity is dependent on error-prone virus replication that is vector species-specific. Previously we demonstrated that WNV genetic diversity is recovered in *Culex* mosquitoes during systemic infection. However, in *Aedes aegypti*, this diversity becomes progressively lower as the virus goes from the bloodmeal to the saliva (11). The degree to which replication in mosquitoes imposes bottlenecks on ZIKV populations have not been described. Repeated bottlenecks are known to reduce virus fitness (14) and therefore represent targets for interrupting virus transmission.

In this study we describe the construction and characterization of a set of ZIKV constructs that contain degenerate nucleotides that result in random “barcodes” within a small region of the genome. Barcoded viruses were characterized *in vitro* and then used to assess bottlenecks in mammalian and mosquito cell lines and in *Ae. aegypti* mosquitoes. We then used an amplicon-based next-generation sequencing technique to quantify the barcode frequencies and measure bottlenecks. These viruses represent a useful tool to study ZIKV population biology within hosts.

## RESULTS

### Barcoded ZIKV viruses replicate at similar levels to wild-type ZIKV

Barcoded viruses have been used to track bottlenecks in different virus systems (13, 15–18). Using an approach similar to Fennessey et al., we inserted genetic barcodes consisting of degenerate nucleotides into the ZIKV genome (strain PRVABC59, GenBank accession number KU501215). Two barcoded viruses were constructed in the coding sequence, and two were constructed in the 3’ UTR (Fig. 1A). In order to do this, we used a novel technique called “bacteria-free cloning” or BFC (Fig. 1B). This technique has the advantage of facilitating the easy propagation of constructs that are toxic to *E. coli* and maintaining the degeneracy of the barcode sequences. Since no bacteria are used, the potential negative effects of lipopolysaccharide during transfection and recovery of virus are also mitigated. Virus replication for all four barcoded viruses and the wild-type clone-derived virus (ZIKV-IC) was assessed on four cell lines. Two mammalian cell lines, Vero and LLC-MK2, were chosen because the former lacks a functional type-1 interferon (IFN) response while the latter does not (19). The two insect cell lines, C6/36 and Aag2, were selected because the former lacks a functional RNAi response and the latter does not (20, 21). Upon infection at a multiplicity of infection (MOI) of 0.01 PFU/cell, all four cell lines supported replication of all four barcoded viruses and the wild-type derived infectious clone (Fig. 2A-D). Very few differences were observed in multi-step growth curves between the viruses. As expected, all viruses replicated faster and to higher peak titers in cells lacking IFN or RNAi responses. Replication in Aag2 cells (Fig. 2D) was poor for all viruses, but titers on days 5 and 6 were significantly higher as compared to day 0 for all viruses except for barcode 2 (bc2), indicative of replication.

**Figure 1.**
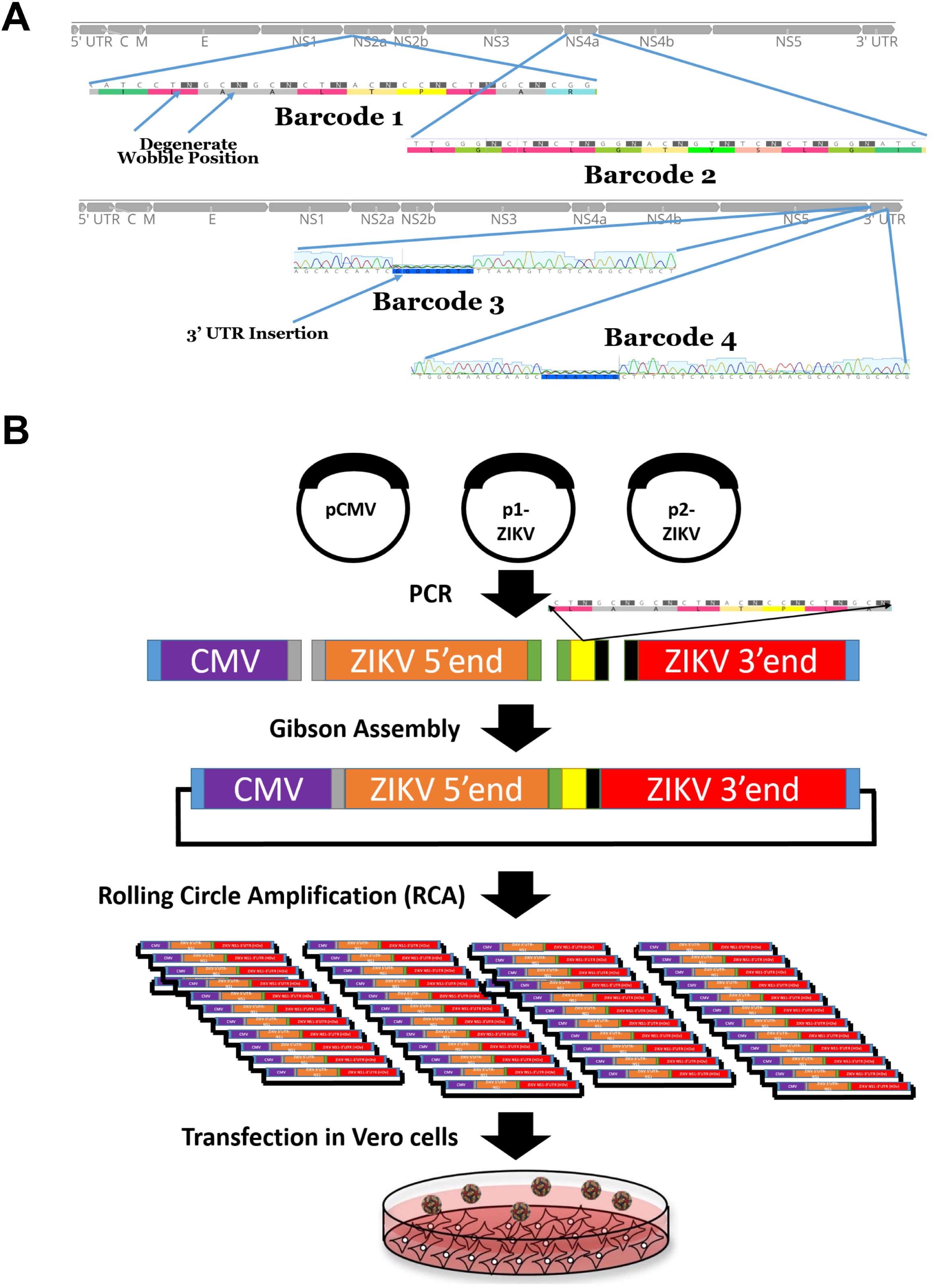
Insertion of genetic barcodes into the ZIKV genome. **A)** Insertion site of degenerate nucleotide barcodes into the genome of ZIKV. Four barcode viruses were constructed, two in the coding sequence and two in the 3’UTR. The coding changes are a consecutive series of degenerate synonymous nucleotide changes at the third codon position. **B)** Schematic of construction of barcode viruses using a bacteria-free cloning (BFC) approach. This approach uses Gibson assembly and rolling circle amplification in place of bacteria. Virus is then rescued by transfection in Vero cells.

**Figure 2.**
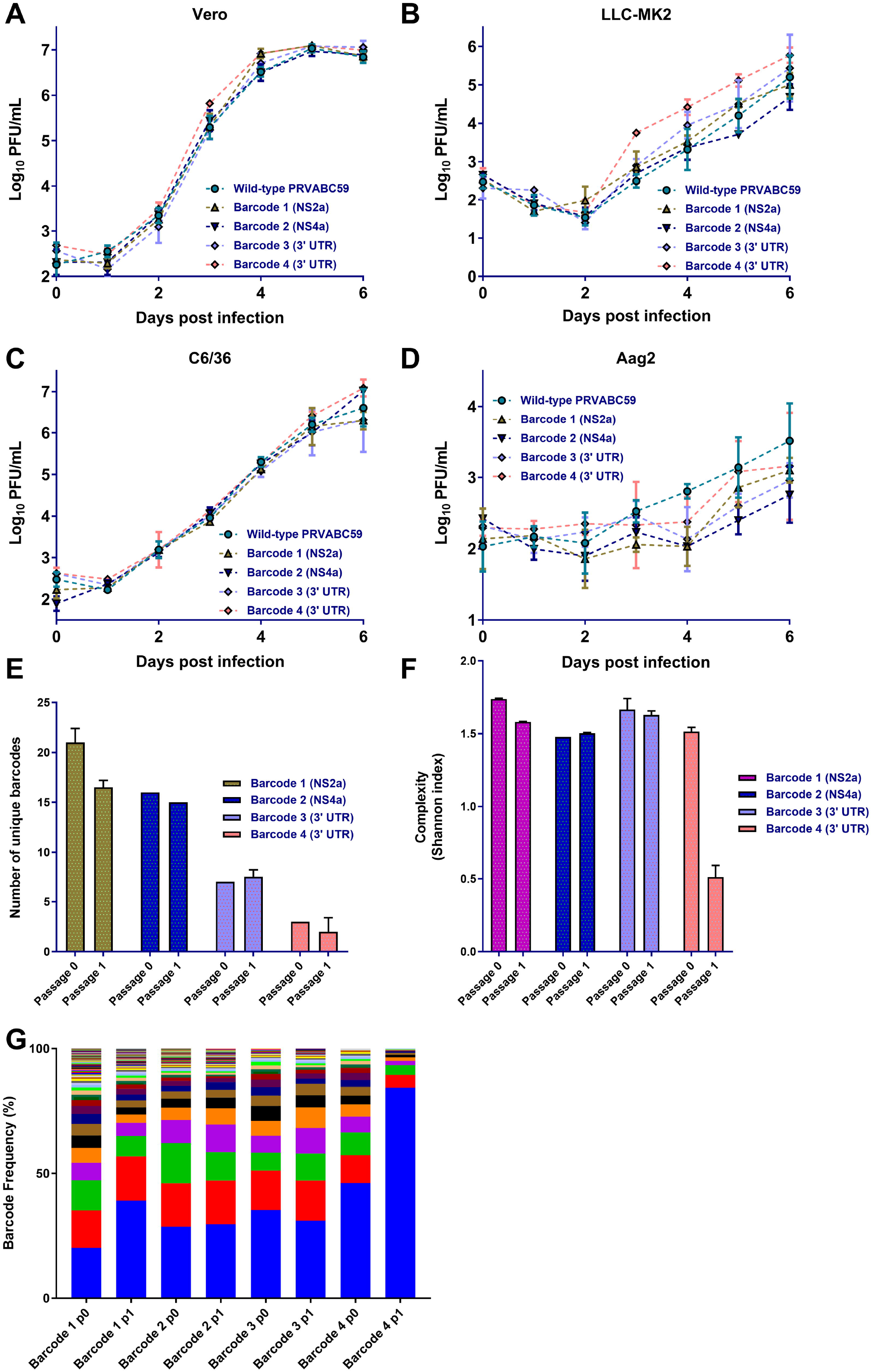
ZIKV barcode viruses replicate similar to wild-type clone derived virus and have differences in barcode diversity. **A-D)** The four barcode viruses have similar replication in two mammalian **(Vero (A) and LLC-MK2 (B))** and two insect-derived **(C6/36 (C) and Aag2 (D))** cell lines. Cells were infected at MOI 0.01. **E**) The number of unique barcodes present in each barcoded virus after transfection (passage 0) or one passage at MOI 0.01 on Vero cells (n=3). **F**) Average genetic complexity at the barcode positions for each barcoded virus. Calculated as – 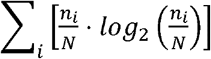. Where *n_i_* is the frequency of each nucleotide at each barcode position and N is the total number of barcode calls at that position **G**) Frequency of individual barcodes. Each color represents a unique barcode. More colors indicate increased barcode diversity.

### Barcodes in the coding sequence are more stable and diverse than those in the 3’ UTR

A high level of nucleotide diversity in the barcode region is necessary to make useful comparisons. Therefore, we sought to characterize the initial barcode composition of all four viruses using next-generation sequencing. By sequencing the wild-type clone stock, which should be predominantly wild-type sequence, we were able to generate a cutoff to differentiate between likely valid and possibly artifactual barcodes present in the samples. The average frequency for the second-highest frequency barcode variant in the wild-type stock ZIKV-IC virus (i.e. the first non-wild-type sequence present) was 0.005. Therefore, we chose this as our cutoff for all barcoded viruses. The number of unique barcodes present in the two coding sequence barcoded viruses (barcoded viruses 1 (bc1) and bc2) was significantly higher than the number of unique barcodes present in viruses with the 3’ UTR insertions (Fig. 2E). This was true in both the transfection-derived stock and after one passage in Vero cells. Diversity in the barcode region was also assessed by calculating the complexity present at the degenerate positions that were inserted in the barcode. In order to do this, we used Shannon’s index, which takes into account the frequency of each nucleotide present in the barcode sequence and the total number of barcode sequences present (22). With greater complexity, the likelihood of encountering the same nucleotide (i.e. the same barcode) becomes lower, indicating greater diversity. Levels of complexity were similar among all barcodes in the initial stock following transfection (Fig. 2F). However, following a passage on Vero cells at MOI 0.01, the barcode 4 (bc4) virus (3’ UTR insertion) had a significant reduction in complexity in the barcode region. The complexity of the other barcoded viruses remained relatively stable following a single passage. The diversity of barcodes present in the population during this passage is presented in Fig. 2G. In the stock viruses derived directly from transfection (passage 0), the diversity of barcoded viruses appears to be greater in bc1 virus than the others, consistent with data in Fig. 2E-F. Given that barcode diversity was highest and replication differences compared to the wild-type infectious clone were lowest in bc1 virus, we chose this for all future studies *in vitro* and *in vivo*.

### Reduced bottlenecks in mosquito cells lacking functional RNAi

*In-vitro* passaging experiments have been instrumental in dissecting basic evolutionary mechanisms and assessing host differences in arboviruses (23–25). Therefore, we used 4 cell lines, Vero, LLC-MK2, C6/36, and Aag2, to analyze bottleneck forces *in vitro* after 3 passages. Due to lower replication rates on LLC-MK2 and Aag2, passaging was performed at MOI 0.001 while an MOI of 0.01 was used on Vero and C6/36 cells. Following the 3 passages, the number of unique barcodes present in each of the samples was reduced significantly in all cell lines except C6/36 (Fig. 3A), which lacks an RNAi response. When compared directly, the number of barcodes in C6/36-passaged virus was significantly higher than in Vero-passaged virus. No differences were observed between LLC-MK2 and Aag2 cells. A similar result was observed in complexity and frequency distributions of barcodes (Fig. 3B-C).

**Figure 3.**
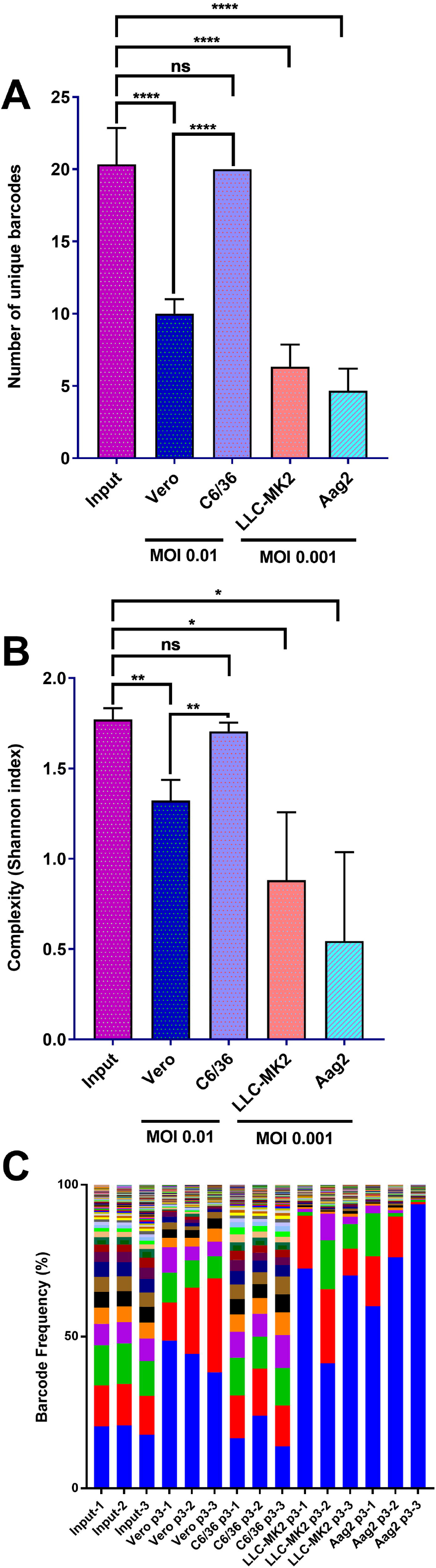
ZIKV barcode viruses undergo cell-type specific reduction in barcode diversity. The four barcode viruses were subjected to 3 three serial passages in two mammalian (**Vero and LLC-MK2**) and two insect-derived (**C6/36 and Aag2**) cell lines. Passages were performed at MOI 0.01 for Vero and C6/36 and 0.001 for LLC-MK2 and Aag2. **A)** The number of unique barcodes present in each barcode virus iteration. **B)** Average genetic complexity at the barcode positions measured by Shannon’s index. Calculated as – 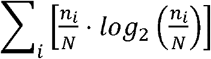. Where *n_i_* is the frequency of each nucleotide at each barcode position and N is the total number of barcode calls at that position **C)** Frequency of individual barcodes. Each color represents a unique barcode. More bars indicate more barcode diversity, n=3 for all treatments. One-way ANOVA was used for all statistical tests.

### A sequential reduction of barcode diversity occurs during infection and dissemination in mosquitoes characterized by stochastic forces

We exposed three different recently field-derived *Ae. aegypti* populations to an infectious bloodmeal containing 1.5e06 PFU/mL of either wild-type clone-derived or bc1 virus. Two of the 3 populations tested (Merida and Poza Rica, herein called Merida and PR) had relatively high rates of infectious virus expectorated in saliva, while one (Coatzacoalcos) had a lower transmission rate (Fig. 4A). Five samples (bloodmeal, midguts, legs, salivary glands (sg) and saliva) were collected from three mosquitoes of each population to represent possible anatomical barriers to virus transmission. For the Merida and PR populations, the drop in barcode complexity and number of unique barcodes from the bloodmeal to the midgut was not significant (Fig. 4B). In contrast, the difference in the number of unique barcodes and the barcode complexity between the bloodmeal and the midgut was significant (p=0.001) and approaching significance (p=0.051, both by one-way ANOVA), respectively for the Coatzacoalcos population. In all populations, both the complexity and number of unique barcodes decreased as the virus spread through the mosquito body to the saliva (Fig. 4B-C). This is clearly visualized in Fig. 4D-F, where in the majority of mosquitoes a single barcode population took over in the legs and predominated through to the saliva. In most cases, the barcode that was predominant in the saliva was different for each individual mosquito tested from each population. Additionally, the highest frequency barcode was not necessarily the highest in the bloodmeal, as even the 7th most abundant barcode in the bloodmeal was able to take over after midgut infection (Fig. 4F). Finally, it was possible for a barcode that was not the most frequent or even absent in the sgs to either take over in the saliva (See Fig. 4, panel E, sample 3) or re-emerge.

**Figure 4.**
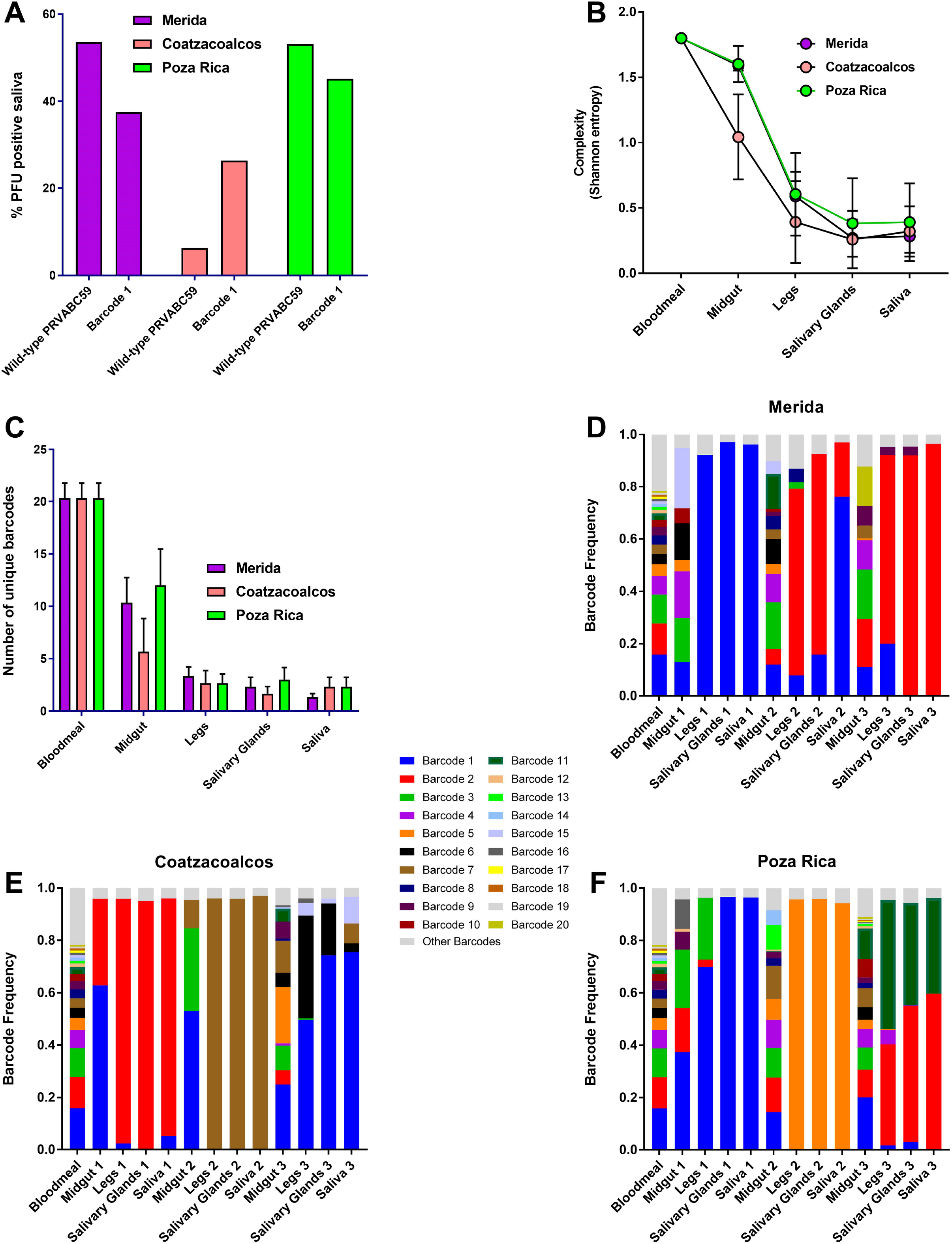
Stochastic forces dominate the sequential reduction in barcode diversity during replication in *Aedes aegypti* mosquitoes. Mosquitoes were given infectious ZIKV bloodmeals and dissected for midguts, legs, salivary glands and saliva 14 days later. **A)** Percent of mosquitoes with infectious virus in the saliva 14 days post-exposure. Comparison between wild-type clone derived ZIKV and barcode 1 virus. n=16-32 for each group. **B)** Average genetic complexity at the barcode positions measured by Shannon’s index. Calculated as – 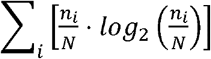. Where *n_i_* is the frequency of each nucleotide at each barcode position and N is the total number of barcode calls at that position. This was measured in the three different mosquito populations across tissue type. **C)** The number of unique barcodes present in each barcode virus iteration. **D-F)** Frequency of individual barcodes from different tissues from the three mosquito populations; Merida **(D),** Coatzacoalcos **(E),** and Poza Rica **(F).** Each color represents a unique barcode. The grey portion represents the remaining proportion of barcodes that weren’t considered “real”. More bars indicate more barcode diversity. n=3 for all treatments. One-way ANOVA was used for all statistical tests.

## DISCUSSION

Population bottlenecks during mosquito transmission have been shown to stochastically reduce viral genetic diversity with significant consequences for viral fitness (11, 13, 26). We therefore developed a barcoded ZIKV to measure population bottlenecks both *in vitro* and *in vivo*. Using a novel approach called bacteria-free cloning we constructed four distinct barcoded viruses and characterized their *in vitro* growth characteristics and barcode composition. Using one of the barcoded viruses, called bc1 virus, we characterized bottlenecks *in vitro* in several cell lines of both mammalian and mosquito origin. C6/36 cells, RNAi deficient *Ae. albopictus* cells, were shown to maintain the most barcode diversity after three passages. In order to characterize bottlenecks *in vivo*, three distinct populations of recently field-derived *Ae. aegypti* mosquitoes were exposed to bc1 virus. Bottleneck strength differed between different mosquito populations of the same species, particularly as the virus infected the midgut. Sequencing of the barcode region in different tissues representing different points for potential bottlenecks emphasized the stochastic nature of replication and transmission of ZIKV in mosquitoes. The barcodes present at the highest frequency in the stock virus commonly were the most abundant in the saliva, although this was not always the case, suggesting that the input virus population structure is a critical determinant of what goes on to be transmitted.

Barcoded viruses have become important tools to investigate virus population dynamics. Several technical strategies have been used to construct these viruses (13, 15–18). While some approaches used a mixture of known tags, more recent approaches have incorporated degenerate nucleotides which increase the theoretical number of barcodes present in a population (18). We sought to use a similar approach and inserted 8 degenerate nucleotides in either the coding sequence (CDS) or 3’UTR of an infectious clone of ZIKV strain PRVABC59, which was originally derived from an infected individual in Puerto Rico in 2015. We found two areas in the region of the genome encoding the non-structural proteins that allowed for the insertion of either 8 or 9 consecutive synonymous changes. For the 3’UTR, we used the RNA secondary structure of ZIKV to identify two regions that would allow insertion of an 8-nucleotide barcode sequence (27).

The flavivirus 3’UTR is highly structured and known to be sensitive to indels that disrupt native RNA folding. Several deletions in the 3’UTR of dengue virus (DENV) have been shown to result in almost complete abrogation of RNA replication in different cell lines (28). In addition, motifs within the 3’UTR are responsible for genome circularization as well as production of sfRNA, both of which are required for efficient replication (29, 30). This may have led to the instability observed in the two barcoded viruses with 3’UTR insertions. While RNA structure in the coding region can be important for function (31–33), those sequences in the two barcoded viruses in the CDS that disrupted this likely would have been quickly removed. Since this was not an insertion, but rather substitutions at 8 nucleotide sites, it is less likely that RNA structure would have had a comparable effect on the number of stable barcodes. It may be that although degenerate nts in the CDS are translationally “silent,” they could impact virus fitness through “translational selection” and codon- and codon-pair biases. Our results on virus replication in various cells and mosquitoes demonstrated that the barcoded viruses replicated at similar levels compared to wild-type clone-derived virus and had similar transmission rates in *Ae. aegypti* mosquitoes, suggesting that the impact of these alterations to the viral genome carried minimal fitness cost. It’s likely that individual barcodes within a population have fitness advantages through codon usage or other mechanisms and certain sequences could have been selected for/against. This is supported by the fact that several barcodes were present at high frequency in the virus stock and further increased in frequency during additional passage. However, the two viruses with barcodes within the coding sequences appeared to maintain higher barcode diversity compared to the 3’UTR insertions, suggesting that selection on the 3’UTR sequence may be stronger than that acting on synonymous changes within the ZIKV CDS. Moreover, these observations allow us to conclude that ZIKV possessing barcodes in the CDS are appropriate tools to measure virus population dynamics.

Using the most diverse barcoded ZIKV, we quantified bottlenecks in several different cell lines *in vitro* and in *Ae. aegypti* mosquitoes *in vivo*. Our results demonstrated that mosquito cells lacking a functional RNAi response place a weak selective pressure on the virus population, especially as compared to RNAi competent cells and even compared to both IFN-deficient and IFN-competent mammalian epithelial cell lines. This is somewhat consistent with previous work using vesicular stomatitis virus (VSV), which was shown to mutate 4x slower in several insect cell lines than in mammalian cell lines (34). In our case this finding appeared to be cell-line-specific, however, as Aag2 cells quickly restricted barcode numbers and complexity at similar levels to LLC-MK2 cells. Therefore, RNAi appears to be a critical component controlling the reduction of virus diversity seen during passaging. It is unclear whether RNAi is selectively reducing specific barcodes due to their sequence or whether the population size is reduced resulting in the loss of lower frequency variants.

We observed differences in bottleneck strengths in different populations of *Ae. aegypti* mosquitoes. This appeared to be related to the transmission rate (TR) of the mosquito population, as the population with the biggest reduction in barcode complexity between the bloodmeal and midgut also had the lowest TR, suggesting the midgut barrier is a major determinant of vector competence. As previously observed with a marked population of VEEV, diversity progressively decreased as the virus passed through anatomical barriers to transmission (13). Previous work showed that WNV haplotypes changed considerably in *Culex* mosquitoes as the virus moved from the midgut to the saliva (11). However, in *Ae. aegypti* mosquitoes, WNV haplotypes remained relatively stable during the same progression. A similar trend was observed here, as the barcodes present in the midgut of one mosquito rarely differed from the virus in the saliva. However, in contrast to that study, we observed a sharp initial drop in virus diversity from the bloodmeal to the midgut and then the legs in infected mosquitoes. This was presumably due to a high amount of complexity (present in the barcoded region) in our starting virus stocks, whereas the cloned virus used in that study had very little initial genetic diversity. Thus, although we started with low overall population diversity on a whole genome level, the barcode sequence provided a high level of starting complexity to assess bottlenecks.

Most of the mosquitoes tested here had one barcode predominate in the legs, salivary glands and saliva, indicating that a severe population bottleneck occurs as virus disseminates from the midgut. However, the virus that predominated in the saliva was not always the highest frequency barcode in the midgut, reflecting the stochastic nature of virus population dynamics, and the importance of genetic drift during mosquito transmission. Visual inspection of the barcode sequences during spread throughout individual mosquitoes (Fig. 4D-F) illustrates this: the vast majority of mosquitoes in each population has a different barcode that predominates in the saliva with significantly reduced total numbers of barcodes and complexity. This result is entirely consistent with previous reports that show that mosquitoes significantly constrain arbovirus evolution (11, 23, 35).

Barcoded ZIKV populations are a promising tool to use for experimental evolution *in vitro* and *in vivo*. These viruses are currently being tested in monkeys (Aliota et al. concomitant submission) and in mice to track viral replication dynamics and bottlenecks in mammalian species. In addition, although the barcoded viruses described here had high complexity, the number of barcodes present did not approach the theoretical limit. With 8 degenerate nucleotides, the theoretical number of barcodes present in a “perfect” stock is ~65,000, considerably higher than what we achieved here. By improving our rescue technique, we have now improved the number of unique barcodes present in the stock virus to a level of almost perfect complexity (data not shown), which will be used in future studies in pregnant monkeys, mice and mosquitoes.

## MATERIALS AND METHODS

### Cells and mosquitoes

Vero (ATCC CCL-81) and LLC-MK2 (ATCC CCL-7) cells were maintained in Dulbecco’s modified Eagle’s medium (DMEM) containing 10% fetal bovine serum (FBS) and 50 μg/mL gentamycin at 37°C with 5% CO_2_. C6/36 cells (ATCC CRL-1660) were maintained in MEM with 10% FBS and 50 μg/mL gentamycin at 28°C with 5% CO_2_. Aag2 cells (obtained from Dr. Aaron Brault) were maintained in Schneider’s insect medium with 10% FBS at 28°C.

*Aedes aegypti* [L.] mosquitoes were collected from wild populations in different parts of Mexico (Merida, Coatzacoalcos and Poza Rica) (Garcia-Luna et al., in submission). The mosquitoes used in this study were between the F2 and F4 generation and were maintained on citrated sheep blood and given 10% sucrose *ad libitum*. Following emergence, adults were maintained under controlled conditions of temperature (28°C), humidity (70% RH), and light (14:10 L:D diurnal cycle). Experiments involving infectious ZIKV in mosquitoes were performed under BSL3 conditions.

### Construction of Barcoded ZIKV

A total of four barcodes consisting of 8 or 9 degenerate nucleotides were constructed, 2 present in the coding sequence at consecutive wobble positions and 2 in the 3’ untranslated region (Figure 1A). Coding-sequence barcodes were selected by searching for consecutive codons in which inserting a degenerate nucleotide in the third position would result in a synonymous change. Barcoded ZIKV was constructed using BFC. First the genome of ZIKV was amplified in two overlapping pieces from the two-part plasmid system previously described (36). The CMV promoter was amplified from pcDNA3.1 (Invitrogen). The barcode region was then introduced in the form of an overlapping PCR-amplified oligo (for the coding sequence) or gBlock (for the 3’ UTR) (IDT, Iowa, USA). Primers used in the study are presented in Table 1. All PCR amplifications were performed with Q5 DNA polymerase (NEB, MA, USA). The amplified pieces were then excised from a gel using crystal violet for DNA visualization (37), thus avoiding the need for exposure to ultraviolet light, and purified with a gel extraction kit (Macherey-Nagel). The purified overlapping pieces were then assembled using Gibson assembly with the HiFi DNA assembly master mix (NEB) and incubated at 50°C for four hours. The Gibson assembly reaction was then digested with Exonuclease I to digest ssDNA, lambda exonuclease to remove non-circular dsDNA, and DpnI to remove any original bacteria derived plasmid DNA at 37°C for 30 minutes followed by heat inactivation at 80°C for 20 minutes. 2 μl of this reaction was then used for rolling circle amplification (RCA) using the Qiagen repli-g mini kit (Qiagen). RCA was performed as instructed except that 2M trehalose was used in place of water for the reaction mix to reduce secondary amplification products (38). Reactions were incubated at 30°C for 4 hours and then inactivated at 65°C for 3 minutes. Correct banding pattern was confirmed by restriction digestion and sequence was confirmed via Sanger sequencing. A schematic depicting the construction and rescue of the barcoded viruses can be seen in Figure 1B.

### Rescue of Infectious Clones

Following completion, the RCA reactions were digested with NruI at 37°C for 1 hour in order to linearize the product and remove the branched structure of the RCA reaction. Generation of an authentic 3’UTR was assured due to the presence of the hepatitis-delta ribozyme immediately following the viral genome. The digested RCA reaction was then purified using a PCR purification kit (Macherey-Nagel) and eluted in molecular grade water. Purified and digested RCAs were transfected into 80-90% confluent T75 flasks of Vero cells using Xfect transfection reagent (Clontech) following the recommended protocol. Infectious virus was harvested upon seeing 50-75% cytopathic effect (CPE), which was 6 days-post transfection. Viral supernatant was then clarified and supplemented to a final concentration of 20% fetal bovine serum (FBS) and 10 mM HEPES before being frozen in singleuse aliquots. The titer was measured by plaque assay on Vero cells.

### *In vitro* replication

Multi-step growth curves were performed on Vero, LLC-MK2, C6/36, and Aag2 cells at an MOI of 0.01. The day before infection, cells were seeded in 12-well plates. Following infection, the virus was allowed to absorb for two hours, at which point virus inoculum was removed, cells were washed with PBS and fresh culture media was added. To monitor replication dynamics, a 100 μL supernatant was harvested and frozen for later titration every day (including day 0) for a total of 6 days. Cells were then supplemented with 100 μL fresh media at each time point. Viral titers were determined by plaque assay on Vero cells.

### *In vitro* passaging experiments

Passaging experiments were performed on Vero, LLC-MK2, C6/36, and Aag2 cells in 12-well plates. Passages on Vero and C6/36 cells were performed at a MOI of 0.01. Due to low replication levels, passages were performed at a MOI of 0.001 in LLC-MK2 and Aag2 cells. Cell culture supernatant was harvested at day 3 post-infection for Vero and C6/36 and day 6 post-infection for LLC-MK2 and Aag2. Infectious virus was then quantified by plaque assay on Vero cells and used to initiate another passage on the corresponding cell line. The passage 3 supernatant was used for NGS to analyze barcode populations.

### Vector competence studies

To assess bottlenecks *in vivo*, adult females from the 3 different populations were exposed to an infectious bloodmeal containing 1.5e06 PFU/mL of either ZIKV wild-type clone virus or barcoded virus. Following each infectious bloodmeal, back-titration was performed to ensure that virus titers were comparable. On day 14, mosquitoes were dissected for midguts, legs/wings, salivary glands and saliva in the same manner as we have previously described (39). Presence of virus was assessed in the saliva using both qRT-PCR (40) and plaque assay.

### Library Prep and Data Analysis

RNA was extracted from all samples using the Mag-Bind Viral DNA/RNA 96 kit (Omega Bio-Tek) on the KingFisher Flex Magnetic Particle Processor (Thermo Fisher Scientific). RNA was eluted in 30□μl nuclease-free water. Regions around the barcode were amplified using primers tagged with Illumina compatible adapters using the NEBNext Ultra II Q5 Master Mix (NEB) adapted for qPCR by adding 5 μM Syto9 green fluorescent nucleic acid stain. Amplicons were then purified with AMPureXP beads (Beckman-Coulter) at a 1.0x ratio. A second indexing PCR was performed using homemade Illumina indexing primers in the same manner as the first PCR, followed by a purification. Libraries were then pooled by volume and sequenced following manufacturer’s instructions on the Nextseq 500 (Illumina) via paired-end sequencing.

Sequencing data was demultiplexed, and fastq files were trimmed of adapter and index sequences. The paired-end reads were then merged using BBMerge, aligned to the ZIKV genome using BBMap, trimmed to the barcode region using Reformat.sh, and barcode sequences were counted and provided in fasta format using kmercountexact.sh. All of the programs listed are a part of the BBTools suite of software (Brian Bushnell, sourceforge.net/projects/bbmap/). The cutoff for barcodes was determined by sequencing the wild-type clone-derived virus around the barcode area. As expected, the wild-type sequence was the predominant sequence present in these sequences (frequency from .96-.99), and the average frequency of the second highest barcode (in this case the first cutoff sequence) was 0.005. This was used as the cutoff for all further analyses. The number of unique barcodes was defined as the number of barcodes called at any frequency higher than the cutoff. The percent of unique barcodes is defined as the number of unique barcodes divided by the total number of called barcode sequences. Genetic complexity was calculated using Shannon’s index using the following equation -

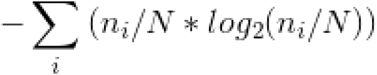

Where n_i_ is the frequency of each nucleotide at each barcode position and N is the total number of barcode calls at that position. A perfectly complex viral population (a barcode sequence with 25% of each nucleotide) would equal 2.

### Statistical analysis

Comparisons of virus titers in growth curves were performed using two-way ANOVA with Tukey’s correction at each time point. One-way ANOVA was used for all other data sets. GraphPad Prism 7.0 (La Jolla, CA) was used for all statistical tests, and significance was defined as p<0.05.

## ACKNOWLEDGEMENTS

We gratefully thank Brian Geiss for discussions involving infectious clones. We would also like to thank Jeff Kieft and Benjamin Akiyama for insight into potential insertion sites into the 3’ UTR.

